# Differential activation of NF-κB and HIF-1α between alveolar-like macrophages and myeloid-derived macrophages drive inflammatory differences following *Mycobacterium abscessus* infection

**DOI:** 10.1101/2025.04.03.647026

**Authors:** Haleigh N. Gilliland, Soledad Soverina, Kayla N. Conner, Taryn E. Vielma, Andrew J. Olive

## Abstract

Pulmonary infections caused by *Mycobacterium abscessus* (Mab), a rapidly growing nontuberculous mycobacterium (NTM), are on the rise in patients with chronic or acquired lung disease. In contrast to immunocompetent individuals, these patient cohorts exhibit abnormal pulmonary function that result from chronic inflammation and mucus build-up. Treatment regimens rely on multi-drug cocktails yet Mab’s natural recalcitrance to common antibiotics extends treatment timelines and increases the frequency of treatment failures. Thus, it is important to understand the mechanisms by which immunocompetent individuals clear Mab with relative ease while susceptible individuals do not. In the lungs, macrophages are the first immune cell Mab encounters following infection, with both resident alveolar macrophages and recruited myeloid derived macrophages playing important roles during infection control. However, the specific role of these distinct macrophage populations in regulating control and inflammatory responses during Mab remains limited due to a lack of *ex vivo* models that recapitulate the functions of different macrophage subsets. Here, we leverage a fetal-liver derived alveolar macrophage (FLAM) model to define early inflammatory responses occurring at the Mab-macrophage interface compared to bone-marrow derived macrophages (BMDMs). Even though both FLAMs and BMDMs similarly control intracellular Mab, the inflammatory response between these macrophage populations is significantly different. BMDMs robustly activated NF-κB transcriptional targets that include important chemokines and inflammatory cytokines like TNF, FLAMs transiently induced these genes following Mab infection. While activation of FLAMs or BMDMs with IFNγ prior to Mab infection did not alter Mab intracellular dynamics, it did drive FLAMs to be more inflammatory, yet important differences remained compared to BMDMs, including the lower expression of the inducible nitric oxide synthase. This was reversed with chemical activation of HIF1α. We conclude that FLAMs and BMDMs differentially respond to Mab infection due to differences in signaling networks activated following innate immune sensing, with FLAMs being more hypoinflammatory than BMDMs. More broadly our results highlight a need to better understand the initial interactions with Mab and distinct macrophage populations to define pathways that contribute to pulmonary protection or disease.

## INTRODUCTION

Pulmonary infections caused by rapidly growing nontuberculous mycobacterium (NTM) are on the rise, with a 400% increase in prevalence from 1987 to 2015 [1–3]. Of these infections, those caused by *Mycobacterium abscessus* (Mab) are the second most common [4]. While the majority of immunocompetent hosts control Mab infections, patients with pre-existing pulmonary conditions including cystic fibrosis (CF), chronic obstructive pulmonary disease (COPD), or bronchiectasis are at a particularly high-risk for chronic Mab pulmonary infection [5–8]. Complicating matters, Mab remains intrinsically resistant to many antibiotics, requiring multi-drug treatment regimens and high rates of treatment failure [9–11]. The lungs of susceptible patient cohorts are characterized by structural damage, inflammation, and/or changes in mucus regulation, changing the pulmonary environment in dramatic ways [7, 12, 13]. How these structural and inflammatory changes directly alter Mab-host interactions remains unclear. With no protective vaccine and limited drug options, understanding how immunocompetent hosts control Mab is of critical importance to develop more clinically relevant host-directed therapies.

Central to host-pathogen interactions in the lungs are macrophages, key innate immune cells tasked with sensing the environment, initiating inflammation, and controlling infections [14, 15]. In the lungs, several macrophage sub-populations including fetal-derived resident macrophages and myeloid-derived interstitial/recruited macrophages play important roles in maintaining pulmonary homeostasis and protecting the lungs against respiratory pathogens [15–17]. The difference in ontogeny and the local environment drives important phenotypic differences in these distinct macrophage subtypes [14, 16]. Resident lung macrophages, such as alveolar macrophages (AMs), are maintained in the airspace to recycle surfactants produced by lung epithelial cells [15–17]. AMs are the first immune cells to detect inhaled pathogens and control initial immune responses during pulmonary infection. Work examining other respiratory infections, including *Mycobacterium tuberculosis*, suggest AMs are hypo-inflammatory and restrain their interactions with T cells to prevent robust adaptive immune activation [18, 19]. In contrast, myeloid-derived macrophages are highly inflammatory cells that drive increased pathogen control and T cell activation [18, 20, 21]. Combined, these inflammatory responses result in tissue inflammation that modulates macrophage function.

During lung infections, inflammatory signals can further modify the local environment to drive protection or pathology [22–24]. One key inflammatory cue is the production of the cytokine interferon-gamma (IFNγ) [25]. This protective cytokine is primarily produced by NK and Th1-activated T cells and is required for humans to control mycobacterial infections, as the loss of IFNγ signaling results in mendelian susceptibility to mycobacterial disease (MSMD) [26, 27]. IFNγ contributes to immune control by upregulating antimicrobial and T cell mediated restriction pathways [19, 28]. These pathways include the production of reactive oxygen species (ROS) and nitric oxide (NO) that directly control pathogens and inflammatory signaling [27, 28]. Recent work suggests IFNγ responses are distinct in different macrophage subtypes, with these differences driving alternative immune functions [18, 29]. Given patients that are susceptible to Mab are characterized by chronic inflammatory lung environments, it is important to understand how distinct macrophage subsets respond to Mab in both resting and inflammatory states.

To date, the majority of studies examining Mab-Macrophage interactions have used murine bone marrow-derived macrophages (BMDMs) and/or immortalized macrophage cell lines including RAW264.7, J774, and Thp1 cells [30–33]. These studies identified important host genes required for the uptake of Mab as well as roles for TLR2 and Nod2 in activating pathways that drive TNF, type I IFN, and NO production to control Mab [30, 34, 35]. Mab infected Zebrafish embryo studies have found TNF signaling and IL8-mediated neutrophil recruitment are required for granuloma formation [36, 37]. However, we continue to lack an understanding of the early interactions between Mab and AMs. One reason for this gap in knowledge is the limited approaches available to understand AMs. AMs are particularly challenging to isolate and to maintain in the AM-like states observed in the lungs. Recent advances in *ex vivo* culturing now enable a mechanistic understanding of interactions with AM-like cells [38–40]. One model we recently developed are fetal liver-derived alveolar-like macrophages (FLAMs) [38]. These cells are isolated from the fetal liver, which is the source of AMs *in vivo*, and cultured with two key lung cytokines GM-CSF and TGFβ [38, 41, 42]. FLAMs are transcriptionally similar to AMs yet distinct from myeloid-derived BMDMs in both resting and IFNγ-activated states [18], thus, serving as a useful model to understand mechanistic AM responses during infection.

Here, we defined macrophage subset specific, early host responses to Mab infection. We first compared the uptake and bacterial growth kinetics in BMDMs and FLAMs to model distinct populations in the lungs in both resting and IFNγ-activated cells finding few differences in Mab control or cell death early following infection. In contrast, we observed significant differences in the innate response in resting and IFNγ-activated FLAMs and BMDMs that was dependent on Mab infection. Our mechanistic studies found differences in the activity of key transcription factors including NF-κΒ and HIF1α that we found partially mediate the differential inflammatory response between FLAMs and BMDMs. These data uncover key differences in the transcriptional response of distinct macrophage subtypes, identifying cell-type specific inflammation that may play an important role regulating disease control or progression during respiratory infections.

## RESULTS

### Mab persists in BMDMs and FLAMs over 48 hours of infection

As a first step to understand differences in the early host-Mab interactions between macrophage subsets we compared whether BMDMs or FLAMs phagocytose Mab with different efficiency. To test this, we infected resting BMDMs derived from HoxB8-mediated conditionally immortalized progenitors or FLAMs with the smooth ATCC Mab strain 19977 expressing constitutively active mEmerald GFP at increasing multiplicities of infection (MOIs) [31]. Four hours later, flow cytometry was used to quantify the percent of cells that were GFP positive. While we noted an increase in the percent of infected cells as the MOI increased, we did not observe any significant differences in uptake between BMDMs and FLAMs (Figure 1A). In addition to percent of infected cells, we wondered whether the mean fluorescence intensity (MFI) of infected cells could be used a surrogate for intracellular bacterial level. To test this prediction, we first infected immortalized BMDMs with mEmerald-Mab and used cell sorting to isolate infected cells with high or low GFP MFI and plated for CFU (Figure 1B). We found that cells sorted from the high GFP MFI contained more Mab on a per cell basis than the same number of cells from the low GFP group (Figure 1C). These data show that MFI is a useful correlate of intracellular bacterial levels. When we examined the MFI of cells infected with increasing MOI of Mab at 4 hours, we observed no significant differences between BMDMs and FLAMs (Figure 1D). These data show that the uptake of Mab by BMDMs and FLAMs is similar.

**Figure 1.**
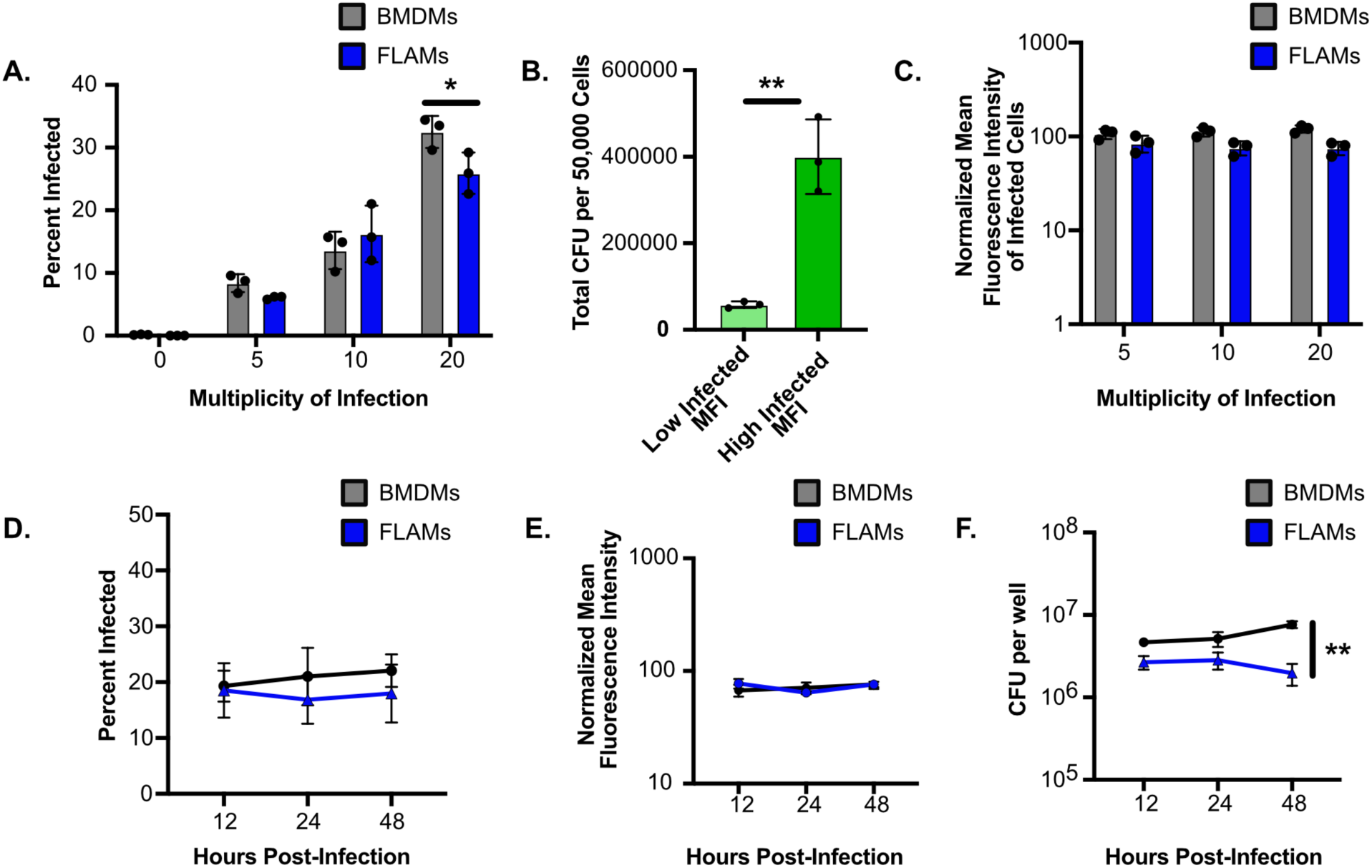
FLAMs and BMDMs similarly take up and control intracellular Mab. **(A)** BMDMs or FLAMs were infected for 4 hours with the indicated MOI of mEmerald-Mab and flow cytometry was used to quantify the percent of cells infected. **(B)** iBMDMs were infected with mEmerald-Mab for three days and 50,000 infected cells with either high mEmerald mean fluorescence intensity (MFI) or low MFI were sorted, plated and total CFU from each population was quantified. **(C)** BMDMs or FLAMs were infected for 4 hours with the indicated MOI of mEmerald-Mab and flow cytometry was used to quantify the MFI of infected cells. **(D)** BMDMs or FLAMs were infected for 4 hours with mEmerald-Mab (MOI 5) and flow cytometry was conducted at the indicated times post-infection to quantify the percent of cells infected and **(E)** the MFI of infected cells from each condition. **(F)** In parallel cells were lysed and intracellular Mab was quantified by CFU plating. Each is experiment is representative of at least 3 independent experiments with at least three biological replicates per experiment. * p<.05 **p<.01 by two-way ANOVA with a Tukey correction for multiple comparisons (A and F) and unpaired t-test for B. All significant comparisons are indicated, and remaining comparisons are not significant.

We next characterized the intracellular dynamics of Mab in distinct macrophage subsets over time. BMDMs or FLAMs were infected with mEmerald-Mab at an MOI of 5 and 12, 24 and 48 hours later flow cytometry was used to monitor intracellular Mab. We observed no significant changes in the percent of infected cells or the MFI of infected cells over this time (Figure 1D and 1E). To confirm these results this experiment was repeated, and cells were lysed for quantification of colony forming units (CFU) (Figure 1F). We found no significant differences in intracellular Mab between cell types over the first 24 hours of infection. However, at 48 hours we observed greater than a three-fold difference between BMDMs and FLAMs that was driven by both a decrease in CFU in FLAMs and an increase in CFU in BMDMs. These data suggest that Mab persists in resting BMDMs and FLAMs over several days with identical intracellular burdens 24 hours following infection.

### Mab infection of FLAMs results in distinct cytokine production independent of cell death

Given the similar intracellular Mab dynamics between BMDMs and FLAMs, we next tested whether there are differences in cell death and inflammation during infection. FLAMs and BMDMs derived from HoxB8-mediated conditionally immortalized progenitors were infected with Mab at an MOI of 5 and 48 hours later cell death was quantified by flow cytometry using a viability dye (Figure 2A). Our results found limited induction of cell death following infection in either cell type using flow cytometry. To confirm these results, we next quantified total ATP via a CellTiter Glo assay as an orthologous cell death approach (Figure 2B). While BMDMs showed no change in cell death we observed a small but significant 10% decrease in the viability of FLAMs following Mab infection at both 24- and 48-hours following infection. These data suggest that over the first two days of infection, when the intracellular levels of Mab are similar between BMDMs and FLAMs, there are no major changes in cell viability.

**Figure 2.**
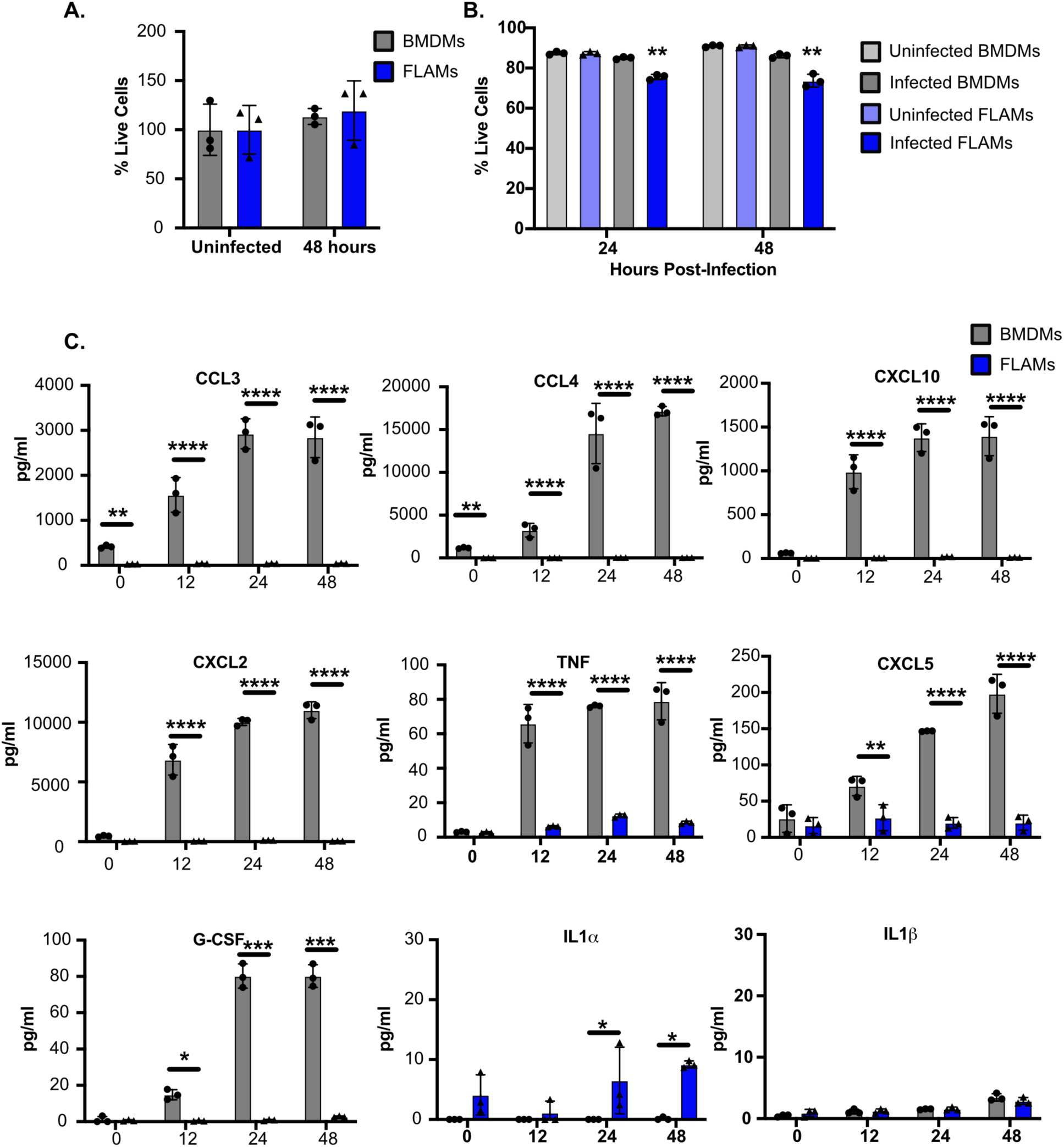
BMDMs are more inflammatory than FLAMs in response to Mab infection without driving more cell death. **(A)** Shown is the percent of viable cells (Live/Dead stain negative) quantified by flow cytometry for BMDMs or FLAMs infected with mEmerald-Mab (MOI 5) for 48 hours. **(B)** The percent of viable cells 24 and 48 hours following mEmerald-Mab infection of BMDMs or FLAMs (MOI 5) using CellTiter Glo. The percent cells that were alive was determined by normalizing each sample to the mean of the uninfected cell type control. **(C)** Shown is the concentration of cytokines from the supernatants of BMDMs or FLAMs infected with mEmerald-Mab (MOI 5) at the indicated timepoints. A & B are representative of three independent experiments with three biological replicates per group. Panel C is from a single multiplex experiment with three biological replicates per group. * p<.05 **p<.01 ***p<.001 ****p<.0001 by two-way ANOVA with a Tukey correction for multiple comparisons.

We next examined whether the inflammatory response was different between Mab infected BMDMs and FLAMs. Cells were infected with Mab at an MOI of 5 and 12, 24 and 48 hours later pro-inflammatory cytokines was quantified in the cell culture supernatants by a multiplex Luminex assay. We observed significant differences between BMDMs and FLAMs, with BMDMs producing 100-10,000 times the levels of the cytokines CCL3, CCL4, CXCL2, CXCL5, CXCL10, TNF and G-CSF compared to FLAMs (Figure 2C). We noted a small but significant 5-fold induction of IL1α only in FLAMs infected for 24 and 48 hours, however, we observed no IL1β produced by either macrophage subset. Taken together, these results show that Mab infected BMDMs have significantly higher and broader inflammatory responses that are independent of changes in bacterial level or cell death.

### Transcriptional analysis finds Mab infection of BMDMs is more inflammatory than FLAMs

We next characterized changes to the global transcriptome of both FLAMs and primary BMDMs over time during Mab infection. Cells were infected with Mab at an MOI of 5, then 6 and 24 hours later RNA was isolated and global mRNA sequencing analysis was performed (Supplementary Table 1). PCA analysis showed a strong separation between BMDMs and FLAMs in all conditions along PC1, in line with our previous studies (Figure 3A) [18]. While there was a major shift in BMDMs at 24 hours following infection along PC2, this was not observed in FLAMs. This suggests that Mab infection induces a less dramatic effect on the transcriptome of FLAMs compared to BMDMs. We next compared genes that were differentially expressed in either FLAMs or BMDMs over time (Figure 3B). We noted 329 and 119 genes were induced or repressed respectively in both FLAMs and BMDMs at all timepoints. We observed that there were 4 times more uniquely induced or repressed genes in BMDMs at 6 hours post-infection compared to FLAMs, a trend that remained at 24 hours post-infection. Pathway analysis of these gene lists found that BMDMs repress oxidative phosphorylation and nucleotide metabolism, while FLAMs induce cell cycle and proliferative genes. To identify other key differences in transcriptional activation during Mab infection of BMDMs and FLAMs, we used clustering analysis to group genes whose expression changes similarly across all conditions. We identified seven unique clusters and conducted pathway and transcription factor analysis to identify transcriptional networks that were associated with each cluster (Figure 3C). Of note, we found that Cluster 3 contained genes that were uniquely induced during Mab infection in FLAMs. KEGG pathway analysis found significant enrichment of genes associated with metabolism, including glycine, serine, and threonine metabolism as well as peroxisomes, a key hub of lipid metabolism. Cluster 5 contained genes that were induced in both FLAMs and BMDMs and was enriched for pathways related to lysosome and phagosomes, as well as antigen presentation. In contrast, Cluster 6 contained genes that were uniquely induced in BMDMs and was enriched for inflammatory pathways including NF-κΒ and TNF. Transcription factor analysis of Cluster 6 agreed with the KEGG pathway analysis and identified a strong NF-κΒ signature. When we more closely examined a subset of NF-κΒ related genes, we found that in BMDMs many of these genes were more highly induced and the expression level of these pro-inflammatory transcripts was higher (Figure 3D). When we examined a subset of chemokines that were elevated in our multiplex analysis above, we noted high, persistent expression of each in BMDMs relative to FLAMs (Figure 3E). The exception to this pattern was IL1α and IL1β. These cytokines were robustly induced at high levels in FLAMs compared to BMDMs, suggesting unique regulation of IL1 in FLAMs. Taken together our transcriptional analysis found that while both FLAMs and BMDMs sense and respond to Mab infection, BMDMs are more inflammatory than FLAMs.

**Figure 3.**
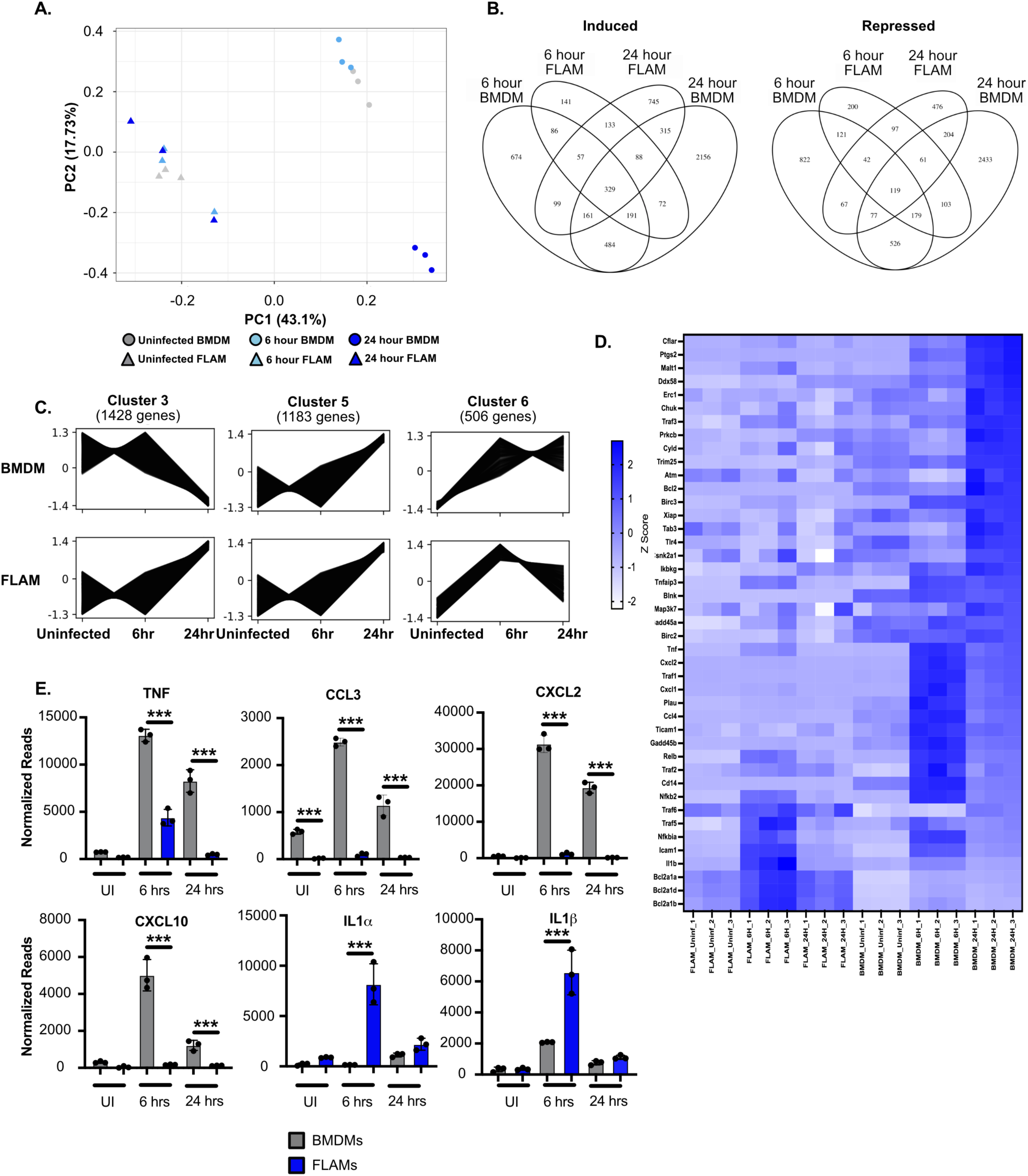
BMDMs induce a stronger NF-κB transcriptional signature during Mab infection compared to FLAMs. **(A)** A principal component analysis (PCA) plot comparing the similarity of the transcriptional responses of BMDMs and FLAMs during Mab infection. **(B)** Venn Diagrams showing shared and unique genes that were significantly induced or repressed in BMDMs and FLAMs during Mab infection using DeSeq2. **(C)** Representative clusters of genes that respond similarly to Mab infection in BMDMs and FLAMs over time. **(D)** Normalized counts of NF-κB genes from the KEGG pathway set were compared across samples of BMDMs and FLAMs infected with Mab over time and is expressed as a heat map. The color scale represents the z-score calculated from normalized read counts across samples for each gene. **(E)** Normalized counts of a subset of NF-κΒ genes that were differentially regulated in BMDMs (gray) and FLAMs (blue) during Mab infection. ***p<.001 based on adjusted *p*-values using DeSeq2 comparisons.

### Differential Nrf2 and NF-**κΒ** activation partially contributes to differences in inflammation of FLAMs and BMDMs during Mab infection

We next examined possible mechanisms contributing to the hypoinflammation we observed in FLAMs following Mab infection. Previous work with *Mycobacterium tuberculosis* infection found that activation of the transcription factor Nrf2 inhibited inflammatory pathways in AMs [43]. To directly test if Nrf2 was responsible for FLAMs hypo-inflammatory response we infected wild type and Nrf2 deficient FLAMs with Mab and examined pro-inflammatory cytokine production over time (Figure 4A). We found that in uninfected resting Nrf2^-/-^ FLAMs there was an increase in the baseline production of TNF compared to wild type cells. Interestingly, we observed no Mab-dependent differences in these cytokines’ infection. These data suggest that while Nrf2 modulates the baseline expression of some inflammatory cytokines in FLAMs, it does not suppress inflammatory signaling in response to Mab infection.

**Figure 4.**
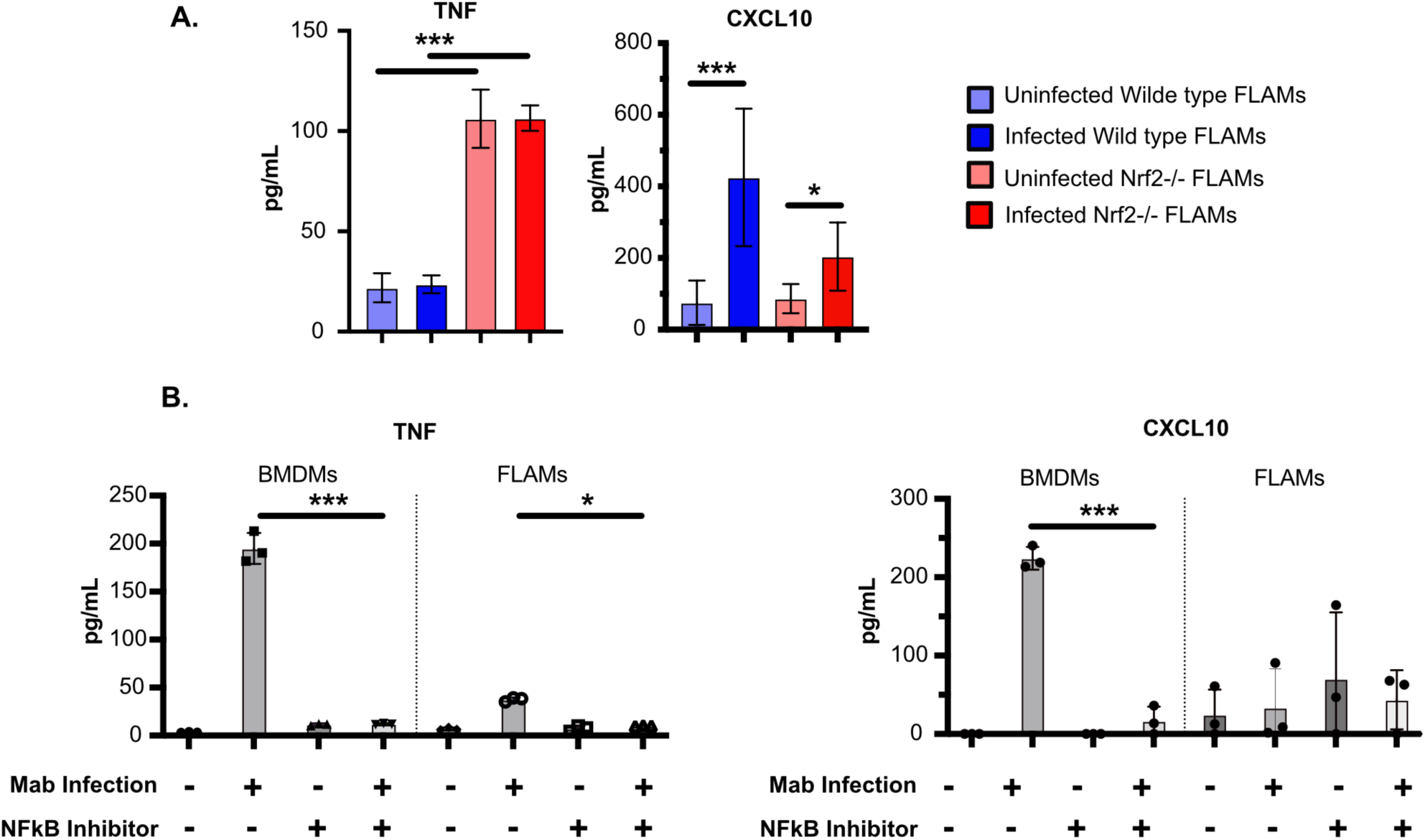
Inhibition of NF-κΒ equalizes the inflammatory response of BMDMs and FLAMs to Mab infection. **(A)** Shown is the concentration of TNF and CXCL10 from the supernatants of wild type or Nrf2^-/-^ FLAMs infected with mEmerald-Mab (MOI 5) for 48 hours by ELISA. **(B)** Shown is the concentration of TNF and CXCL10 from the supernatants of BMDMs or FLAMs infected with mEmerald-Mab (MOI 5) for 48 hours in the presence and absence of the NF-κB inhibitor (5μM). Data are representative of two independent experiments with at least 3 biological replicates. * p<.05 ***p<.001 by two-way ANOVA with a Tukey correction for multiple comparisons. All significant comparisons are indicated, and remaining comparisons are not significant.

Our transcriptional analysis suggested that BMDMs induce NF-κΒ pathways more robustly than FLAMs. To examine the role of NF-κΒ in the hyper-inflammatory response of BMDMs, we inhibited NF-κΒ using the chemical inhibitor BAY 11-7082 in both FLAMs and BMDMs and quantified TNF, an NF-κΒ dependent transcript, by ELISA 6 and 24 hours following Mab infection (Figure 4B) [45]. We observed that blocking NF-κΒ in Mab-infected BMDMs resulted in a 20-fold decrease in TNF a level similar to infected FLAMs. These data suggest that NF-κΒ activation in BMDMs drives higher levels of inflammatory cytokines than FLAMs during Mab infection.

### Activating BMDMs or FLAMs with IFN**γ** does not alter Mab control

Patients who develop chronic Mab respiratory infections are characterized by previous lung damage and ongoing respiratory disfunctions [9, 46]. In these patients, the baseline inflammatory state of macrophages is different from resting macrophages. IFNγ is produced during inflammatory responses and is a key cytokine that activates macrophages and can drive restriction of intracellular pathogens [27, 28]. To test if IFNγ differentially alters Mab-host interaction in FLAMs and BMDMs, cells were activated overnight with IFNγ, then infected with mEmerald-Mab before multiple parameters were examined. First, we examined whether IFNγ altered Mab uptake by BMDMs and FLAMs. We found that four hours following infection, IFNγ did not alter uptake efficiency between BMDMs and FLAMs at increasing MOIs (Figure 5A and 5B). We next examined intracellular dynamics overtime. Using flow cytometry, we noted a slight increase in the percent infected cells in both BMDMs and FLAMs, but this remained stable through 48 hours of infection (Figure 5C). We also noted a similar MFI of infected cells between resting and IFNγ-activated BMDMs and FLAMs (Figure 5D). In line with our flow cytometry approaches, when we quantified viable CFU, we found no significant differences in Mab viability in IFNγ-activated FLAMs or BMDMs (Figure 5E). These data suggest that IFNγ-activation does not significantly alter the intracellular dynamics of Mab, which continues to persist in both activated BMDMs and FLAMs.

**Figure 5.**
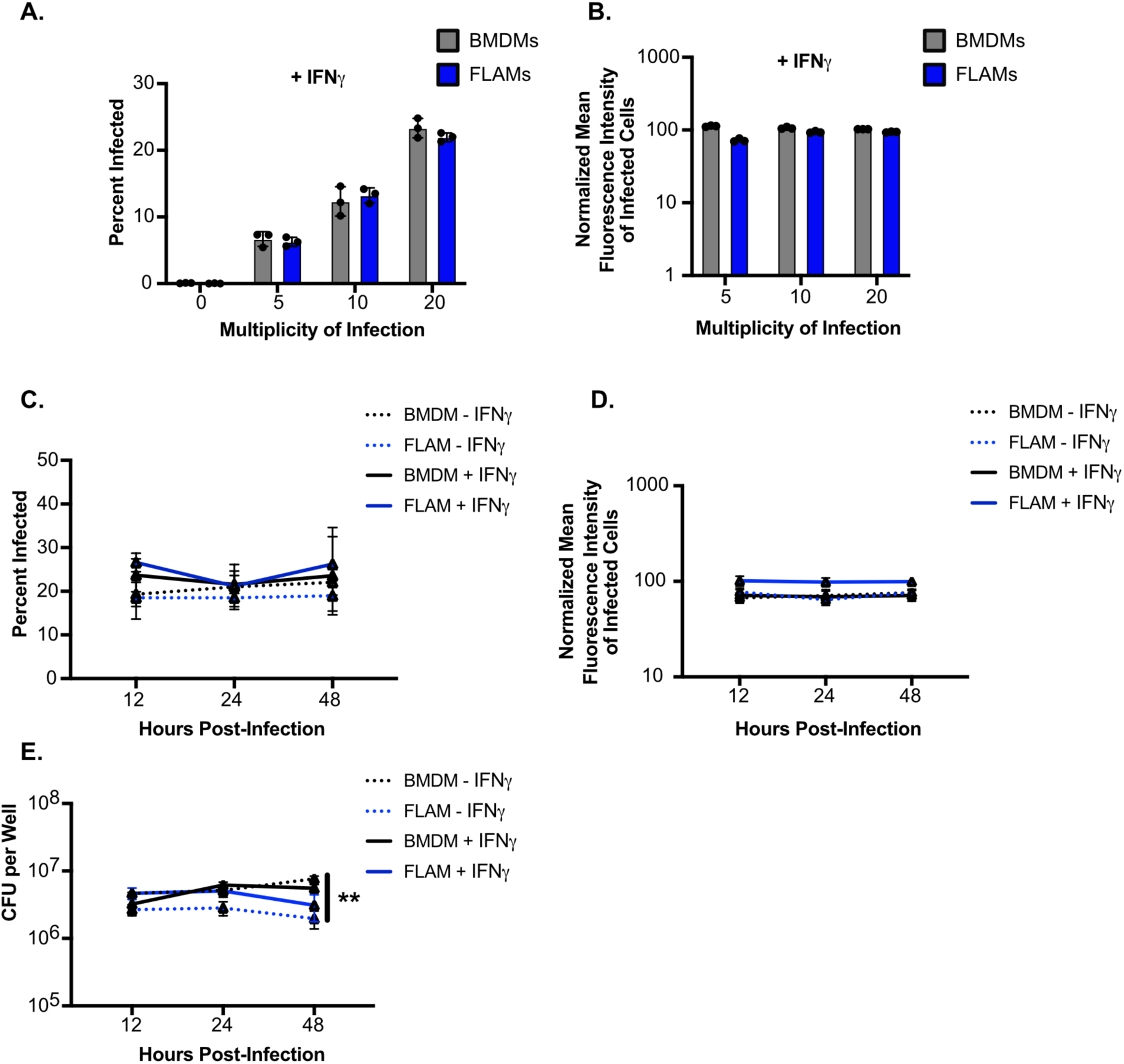
IFNγ activation of BMDMs or FLAMs does not alter Mab uptake or intracellular control. **(A)** The percent of infected cells and the **(B)** MFI of infected cells were quantified by flow cytometry from IFNγ-activated (25ng/ml) BMDMs or FLAMs that were infected with mEmerald-Mab at the indicated MOIs for 4 hours. **(C)** The percent of infected cells and the **(D)** MFI of infected cells were quantified by flow cytometry from untreated or IFNγ-activated (25ng/ml) BMDMs or FLAMs that were infected with mEmerald-Mab (MOI 5) for the indicated times post-infection. **(E)** Viable intracellular Mab was quantified by CFU assay from untreated or IFNγ-activated (25ng/ml) BMDMs or FLAMs that were infected with mEmerald-Mab (MOI 5) for the indicated times post-infection. Shown data are representative of three independent experiments with three replicates per group. Untreated data are from the same experiment shown in Figure 2.1D-2.1E and appropriate multiple hypothesis testing corrections have been made. No significance was observed in panels A and B. For C and D no significance was observed between IFNg-activated BMDM and FLAMs or between resting and activated BMDMs or FLAMs. For E **p<.01 by two-way ANOVA with Tukey correction between IFNg-activated FLAMs and IFNg-activated BMDMs.

### IFN**γ** drives more cell death and inflammatory cytokines following Mab infection in both BMDMs and FLAMs

We next examined how IFNγ alters the host response to Mab infection. IFNγ activated BMDMs and FLAMs were infected with mEmerald Mab and changes in cell viability were quantified over time using two orthologous approaches. First, cell viability was quantified using flow cytometry 12-, 24- and 48-hours post-infection (Figure 6A). Our results showed significant cell death in all conditions independently of infection. In addition, total ATP was quantified using a CellTiter Glo assay (Figure 6B). Our results found that while IFNγ induced significant levels of cell death that approaches 50 percent in both assays, this remained unchanged following Mab infection. These data suggest that while IFNγ activation increases cell death pathways in both BMDMs and FLAMs compared to resting cells, it does so independently of Mab infection.

**Figure 6.**
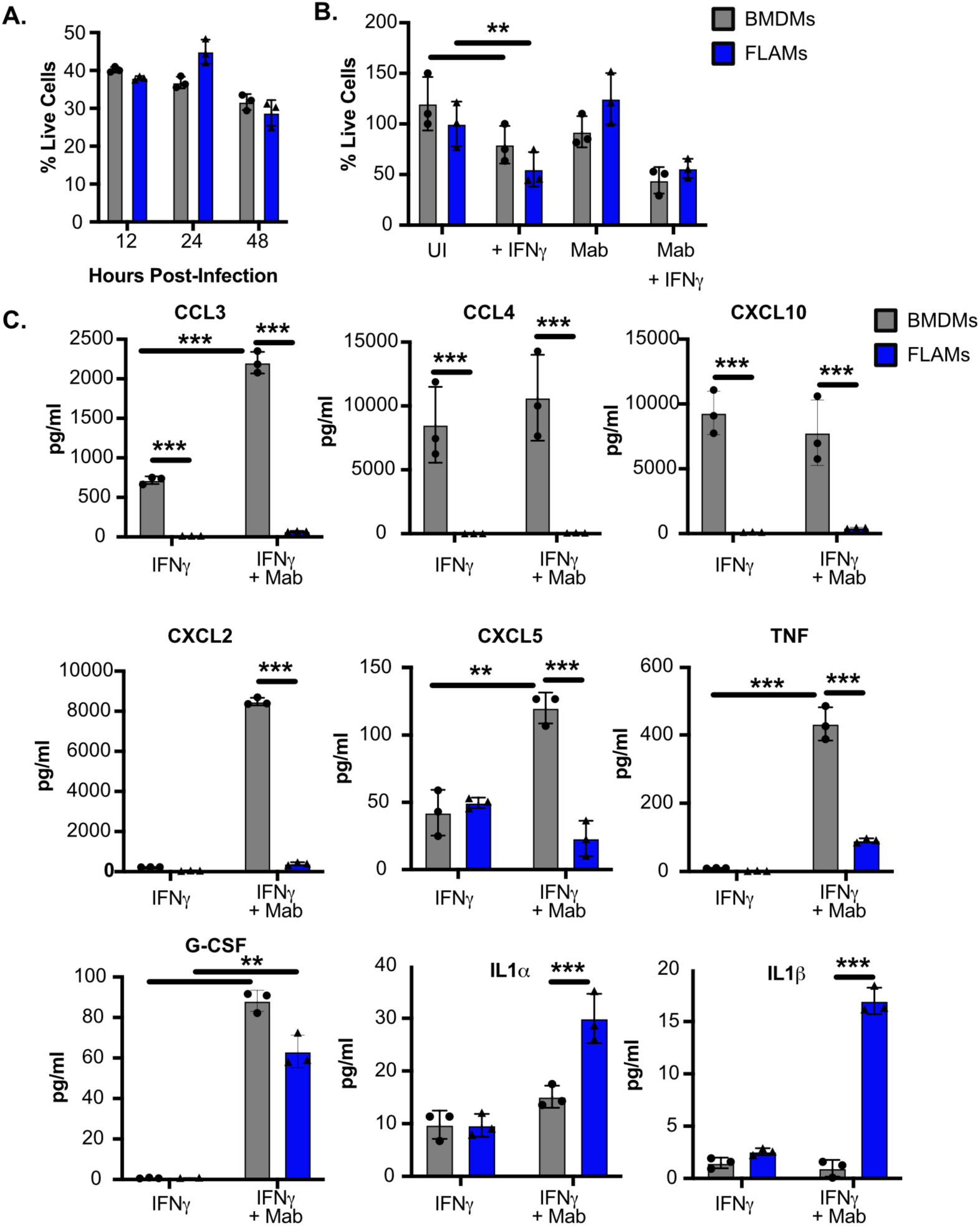
The inflammatory response of IFNγ-activated BMDMs and FLAMs remain distinct. **(A)** Shown is the percent of viable cells (Live/Dead stain negative) quantified by flow cytometry for IFNγ-activated (25ng/ml) BMDMs or FLAMs infected with mEmerald-Mab (MOI 5) for the indicated time. **(B)** The percent of viable cells 48 hours following mEmerald-Mab infection of untreated or IFNγ-activated (25ng/ml) BMDMs or FLAMs (MOI 5) using CellΤiter Glo. The percent cells that were alive was determined by normalizing each sample to the mean of the uninfected cell type control. **(C)** Shown is the concentration of cytokines from the supernatants of IFNγ-activated (25ng/ml) BMDMs or FLAMs infected with mEmerald-Mab (MOI 5) and at the indicated timepoints. A & B are representative of three independent experiments with three biological replicates per group. Panel C is from a single multiplex experiment with three biological replicates per group. **p<.01 ***p<.001 by two-way ANOVA with a Tukey correction for multiple comparisons.

We next dissected if the inflammatory response of BMDMs and FLAMs was altered by IFNγ activation. IFNγ-stimulated FLAMs or BMDMs were infected with Mab at an MOI of 5 and 48 hours later pro-inflammatory cytokines in the supernatants were quantified by a multiplex Luminex assay (Figure 6C). We observed that IFNγ activation alone drove over 100-fold increases off CCL3, CCL4 and CXCL10 in BMDMs but only CCL3 further increased an additional 4-fold following Mab infection. IFNγ-activated FLAMs did not robustly induce these cytokines in any condition. Similar to resting cells, IFNγ-activated BMDMs induced high levels of CXCL2 and CXCL5 following Mab infection, but this was not observed in FLAMs. Notably, G-CSF was similarly induced in both IFNγ-activated BMDMs and FLAMs. We observed that TNF was more robustly induced in IFNγ-activated FLAMs than resting FLAMs, yet the overall TNF levels remained significantly lower than BMDMs. Finally, we found that while both IL1α and IL1β were significantly induced in FLAMs but not BMDMs, the levels of induction were generally low. Together these results show that while IFNγ alters the inflammatory response of both BMDMs and FLAMs, these cell types continue to drive distinct cytokine profiles following Mab infection.

### IFN**γ**-activation drives distinct transcriptional changes in Mab infected FLAMs and BMDMs

We next compared the global transcriptomic changes of IFNγ-activated FLAMs and BMDMs during Mab infection. BMDMs and FLAMs were activated with IFNγ overnight then infected with Mab at an MOI of 5. 6 and 24 hours later RNA was isolated from cells and RNA sequencing analysis was performed. To identify general differences in the transcriptomes, we visualized the data using principal component analysis (Figure 7A). When we examined only IFNγ-activated conditions with BMDMs and FLAMs, we noted separation between cell types along the PC1 axis. We noted separation on PC2 axis for BMDMs infected with Mab for 24 hours. In contrast, we did not observe any significant movement along PC2 with FLAMs. This analysis was remarkably similar to our PCA from resting BMDMs and FLAMs (see Figure 3A). To directly compare resting and IFNγ-activated cells, we generated a new PCA from all conditions (Figure 7A). While FLAMs clustered together for all conditions, both resting and IFNγ-activated BMDMs infected with Mab for 24 hours showed a strong shift from the other BMDMs. These data suggest that Mab infection drives a more robust transcriptional change in BMDMs compared to FLAMs.

**Figure 7.**
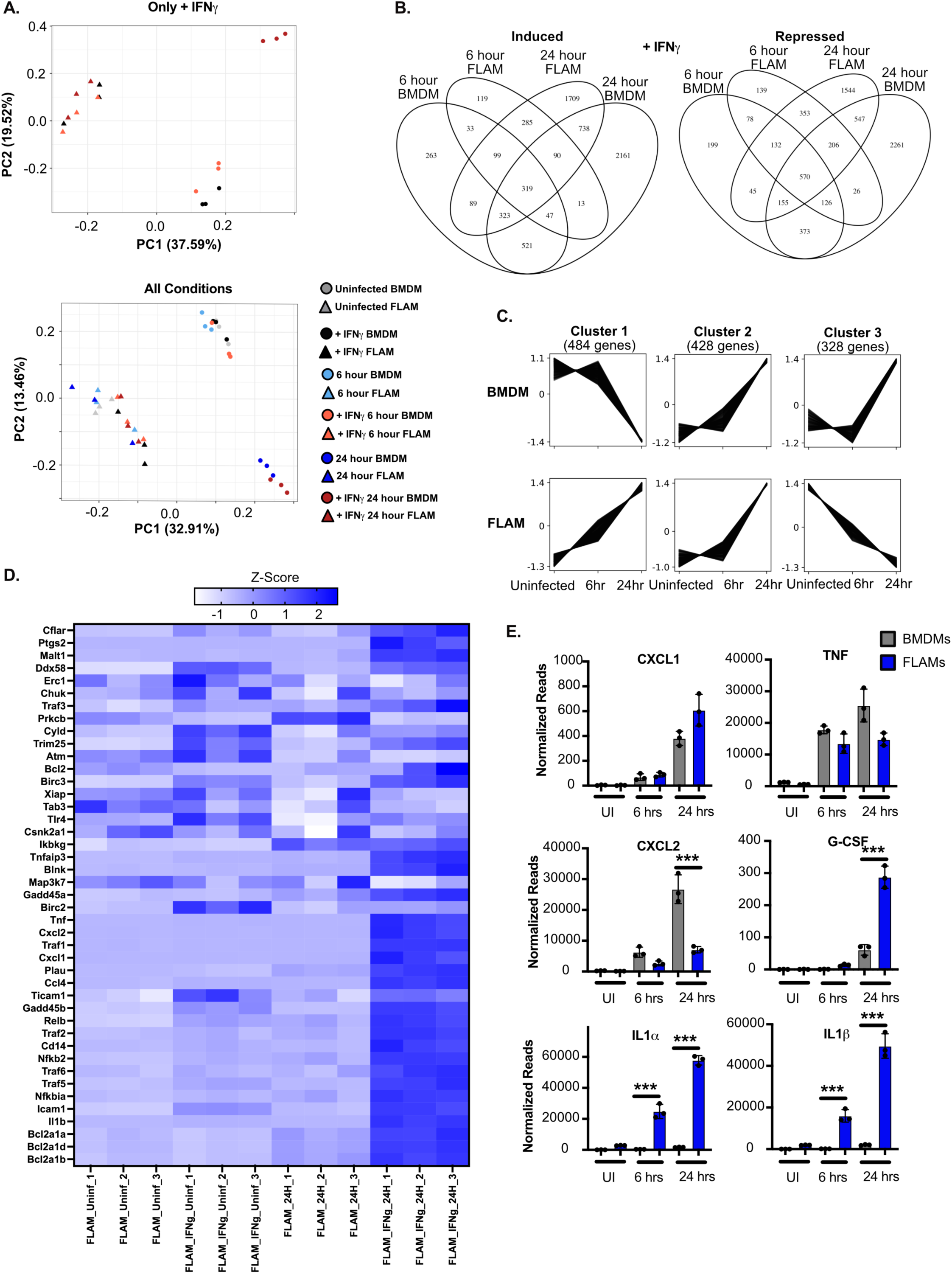
IFNγ drives more robust NF-κΒ activation in Mab infected FLAMs. **(A)** A principal component analysis (PCA) plot comparing the similarity of the transcriptional responses of (Top) IFNγ-activated BMDMs and FLAMs during Mab infection and (Bottom) all conditions examined in BMDMs and FLAMs during Mab infection. **(B)** Venn Diagrams showing shared and unique genes that were significantly induced or repressed in IFNγ-activated BMDMs and FLAMs during Mab infection using DeSeq2. **(C)** Representative clusters of genes that respond similarly to Mab infection in IFNγ-activated BMDMs and FLAMs over time. **(D)** Normalized counts of NF-κΒ genes from the KEGG pathway set were compared across the indicated conditions in FLAMs and is expressed as a heat map. The color scale represents the z-score calculated from normalized read counts across samples for each gene. **(E)** Normalized counts of a subset of NF-κΒ genes that were differentially regulated in BMDMs (gray) and FLAMs (blue) during Mab infection. Statistical significance was determined based on adjusted *p*-values using DeSeq2.

We next compared genes that were differentially expressed in either IFNγ-activated FLAMs or BMDMs over time following Mab infection (Figure 7B). Similar to resting cells we noted almost two times the number of unique induced and repressed genes in BMDMs at both 6 hours and 24 hours post-infection compared to FLAMs. When we compared the lists of differentially expressed genes from IFNγ-activated and resting cells from the same condition, we found that IFNγ drives more changes to gene expression than Mab infection alone in both FLAMs and BMDMs. To identify key patterns of transcriptional changes during Mab infection of BMDMs and FLAMs, we used clustering analysis to identify groups of genes whose expression changes similarly across each IFNγ-activated condition. We found four clusters of genes and using pathway and transcription factor analysis we identified transcriptional networks that were associated with each cluster (Figure 7C). Interestingly, cluster 1 contains genes that are induced in IFNγ-activated Mab infected FLAMs but are repressed in BMDMs. Among the top enriched KEGG pathways in this cluster was oxidative phosphorylation, suggesting key differences in the metabolic response of IFNγ-activated BMDMs and FLAMs to Mab infection. We also observed that Cluster 3 contained genes that were induced in IFNγ-activated Mab infected BMDMs but not FLAMs. In contrast to resting cells, we did not observe a significant enrichment of the NF-κΒ pathway from this cluster. This suggests that IFNγ-activation of FLAMs drives a more robust NF-κB response following Mab infection. To examine this directly we compared the normalized reads from FLAMs for NF-κB dependent genes examined in Figure 3 above. We found that IFNγ-activation of FLAMs drove significantly higher expression of NF-κB dependent genes following Mab infection (Figure 7D). Thus, NF-κΒ can be more robustly activated in Mab infected FLAMs following IFNγ-stimulation. When we examined a subset of genes examined by our multiplex analysis, we saw strong agreement for the majority of cytokines including CXCL1, CXCL2, TNF and G-CSF (Figure 7E). Interestingly, while we also observed that only FLAMs induce IL1α and IL1β in response to Mab, the magnitude of these changes at the transcriptional level was significantly higher compared to the released cytokines.

### The lack of Nos2 expression following Mab Infection of IFN**γ**-activated FLAMs is overcome by the induction of HIF1a

An important role of IFNγ-activation is to induce antimicrobial compounds, such as Nitric Oxide (NO). NO is known to play an important role during Mycobacterial infections, yet if it is differentially regulated during Mab infection remains unclear [21, 47–49]. When we compared gene sets between FLAMs and BMDMs during IFNγ activation, we noted significant differences in the expression of Nos2 and Ptgs2 (Figure 8A). Previous studies found these genes are induced in both a HIF1α-dependent manner that is also associated with a metabolic shift to aerobic glycolysis [50–52]. Given that we observed a specific increase in oxidative phosphorylation pathways in IFNγ-activated FLAMs, we hypothesized that HIF1α may not be as robustly induced in FLAMs compared to BMDMs. When we examined the expression of HIF1α in our RNAseq dataset, we observed that while HIF1α expression was significantly induced during Mab infection of IFNγ-activated BMDMs, the expression remained unchanged in FLAMs (Figure 8B). We next wanted to test whether the activation of HIF1α was sufficient to increase Nos2 expression in FLAMs. FLAMs were left resting or were activated with IFNγ in the presence or absence of the HIF1α activator, DMOG. Cells were infected with Mab and 24 hours later RNA was isolated and the expression of Nos2 was quantified by RT-PCR. We found very IFNγ-activated Mab infected FLAMs treated with DMOG induced 10 times more Nos2 that vehicle control treated cells (Figure 8C). When we repeated this experiment and examined nitrite production using a Griess assay, we observed a similar result (Figure 8D). FLAMs only produced detectable levels of nitrite in the presence of DMOG following IFNγ-activation and Mab infection. These data suggest that HIF1α is not robustly induced in IFNγ-activated FLAMs during Mab infection, and this results in changes to the inflammatory response including low induction of Nos2 and nitric oxide production. Thus, IFNγ-activated FLAMs activate distinct pathways following Mab infection compared to BMDMs.

**Figure 8.**
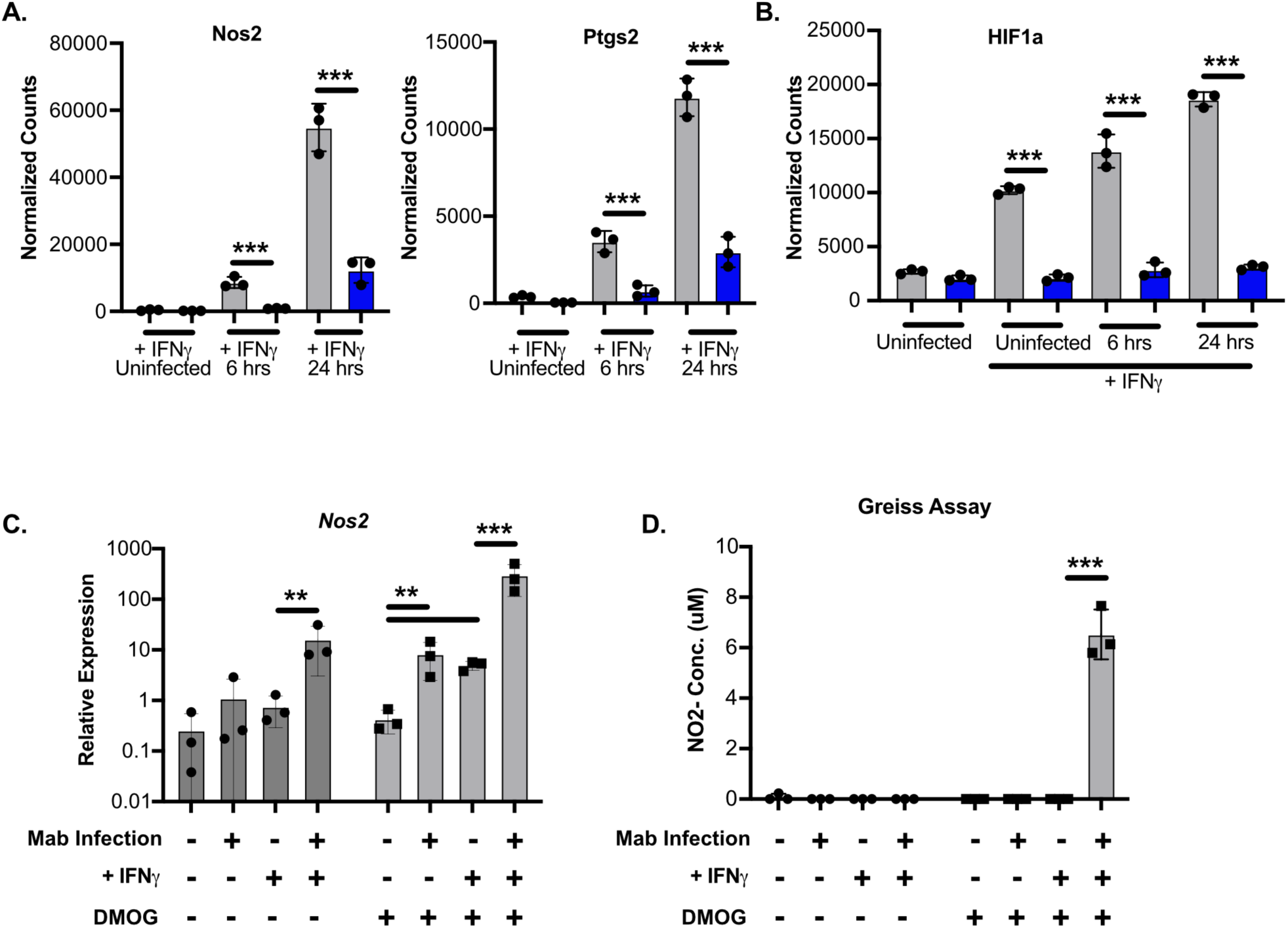
Chemical activation of HIF1α increases Nos2 expression in IFNγ-activated FLAMs following Mab infection. **(A)** Normalized counts of Nos2 and Ptgs2 in IFNγ-activated BMDMs (gray) and FLAMs (blue) during Mab infection. **(B)** Normalized counts of HIF1α in resting or IFNγ-activated BMDMs (gray) and FLAMs (blue) during Mab infection from the RNAseq dataset. **(C)** FLAMs were left untreated or were treated with 25ng/ml of IFNγ and/or 250μM DMOG and 12 hours later cells were infected with Mab for 4 hours. 24 hours later, RNA was isolated, and qRT-PCR was used to determine the relative expression of *Nos2* compared to *B-actin* controls. **(D)** FLAMs were left untreated or were treated with 25ng/ml of IFNγ and/or 250μM DMOG and 12 hours later cells were infected with Mab for 4 hours. 24 hours later, 50μL of cell supernatants were used to measure the concentration of NO2-at each condition. Statistical significance for normalized reads was determined based on adjusted *p*-values using DeSeq2. **p<.01 ***p<.001 by two-way ANOVA with a Tukey correction for multiple comparisons.

## DISCUSSION

Respiratory infections with Mab are an increasing clinical concern. Given the high failures rates of antibiotic therapy and no protective host-directed therapies or vaccines available, understanding how Mab is effectively controlled in some hosts and not in others is critically important. Here, we leveraged new *ex vivo* models of lung-specific alveolar macrophages to better understand key differences in the innate response during Mab infection between distinct macrophage subtypes. Our data suggest that while both BMDMs and FLAMs control Mab infection independently of IFNγ-activation, they sense and respond to Mab infection in distinct ways. Myeloid-derived BMDMs were more inflammatory driving higher levels of cytokines, chemokines, and activated markers, in contrast FLAMs were less inflammatory with lower expression of inflammatory genes. Our findings highlight the importance of defining host-pathogen interactions in a range of tissue relevant immune cells to better identify mechanisms that may contribute to control of Mab and other respiratory infections.

Mab infects macrophages which serve as a key intracellular niche throughout infection [53, 54]. The intracellular dynamics of Mab, however, remain relatively uncharacterized. In contrast to other studies, our findings using an array of readouts including bulk CFU counts as well as single cell flow cytometry approaches suggest that Mab does not replicate to high levels in either BMDMs or FLAMs. Instead, intracellular Mab is relatively steady over the first 2 days of infection while not inducing high levels of cell death. Whether this steady state is indicative of static growth, or equivalent rates of growth and death remains to be determined but will be important to better understand the mechanisms controlling early Mab interactions with macrophages. Given the importance of IFNγ in controlling mycobacterial infections and the inflammatory environment seen in patients susceptible to Mab infection, it was surprising that we found very little effect of IFNγ-activation on intracellular dynamics of Mab infection. This is in contrast to *Mycobacterium tuberculosis* (Mtb), which is readily restricted in IFNγ-activated BMDMs. It is possible that Mab is more resistant than Mtb to intracellular antimicrobial poisons. Alternatively, Mab may be maintained in a distinct intracellular compartment that is resistant to IFNγ-inducible effectors. Future work will be needed to better understand the intracellular dynamics of Mab infections, with particular focus on important macrophage pathways that control the replication of this opportunistic pathogen.

A key finding throughout our study was the distinct inflammatory responses that Mab activated in FLAMs and BMDMs. In general, BMDMs were more hyper-inflammatory, producing higher levels of chemokines like CXLC1 and CXCL2 and cytokines like TNF and G-CSF in both the presence and absence of IFNγ activation. One possible mechanism driving inflammatory differences is the activity of transcriptions factors, such as Nrf2 and NF-κΒ. Our data suggests that while loss of Nrf2 results in increased basal inflammation, it does not alter Mab dependent inflammatory responses in FLAMs. In contrast, we found that NF-κΒ was driving high expression of a subset of cytokines including TNF and CXCL10 in BMDMs and that NF-κΒ was not robustly induced in FLAMs. While IFNγ activation drove more NF-κΒ activation in FLAMs, many key genes remained lower than BMDMs. This included *Nos2*, which we found could be increased in FLAMs treated with IFNγ and the HIF1α activator DMOG. HIF1α is known to be interlinked with the shift of IFNγ-activated cells towards aerobic glycolysis [51, 52]. Thus, taken together we speculate that one key driver of the distinct activation of NF-κΒ and HIF1α in FLAMs and BMDMs during Mab infection is metabolic differences. Understanding how metabolism drives differences between FLAMs and BMDMs and manipulating these pathways genetically or chemically in resting, activated and infected conditions will be an important next step to define the underlying mechanisms of inflammatory regulation in distinct macrophage subsets.

Mab respiratory infections are cleared by the majority of immunocompetent patients, yet the mechanisms mediating this control remain unclear. AMs are the first line of immune defense and our results using FLAMs as a model for AMs suggests these cells control Mab infection without driving a severe inflammatory response. Patients who are susceptible to Mab have pre-existing lung conditions that disrupt the normal state of alveolar macrophages through activation, including IFNγ-stimulation, and the recruitment of myeloid-derived macrophages. The inflammatory environment in the lungs and the ontogeny of airway macrophages during Mab infection may further contribute to inflammation, driving more tissue damage and increasing the severity of respiratory disease. Continuing to dissect how ongoing inflammation in susceptible patients dampens pathogen control without driving lung disease will aid in the identification of new therapeutic opportunities that could be used synergistically with antibiotic therapy.

One important shortcoming of our study is the Mab strain used is a smooth colony variant. It is well known that Mab isolates can present as smooth variants through the expression of GPLs, with the loss of these GPLs resulting in rough colony variants [54–56]. In clinical studies, both smooth and rough variants are isolated from patients, with data suggesting the host environment drives increased transition from the smooth to rough variant during infection [54, 57]. Rough Mab drives more inflammatory cytokines, such as TNF, compared to smooth Mab when infecting BMDMs [53, 58]. Rough isolates are also suggested to result in bacterial cording phenotypes that further drive inflammatory responses [37, 55]. Whether other bacterial factors in addition to GPL expression drive inflammation in macrophages remains to be fully understood. Furthermore, how AMs or FLAMs differentially sense and induce inflammation against rough and smooth Mab isolates and other bacterial factors that modulate host inflammation remains to be tested.

Taken together our study identified many key differences in the inflammatory response of distinct macrophage subsets following infection with Mab. These differences in the innate immune response occurred independently of differences in bacterial control or cell death. Thus, the innate immune wiring of distinct macrophage subsets is unique. These results can now be used to identify the importance of distinct innate sensing pathways during infection with Mab and other respiratory pathogens to develop new host-directed therapies that prevent infection while limiting inflammatory damage to the lungs.

## MATERIALS AND METHODS

### Mab culture conditions

mEmerald GFP-expressing Mab (ATTCC 19977) strains were generated as previously described [31]. All mEmerald GFP-expressing Mab cultures were grown aerobically at 37°C in Middlebrook 7H9 medium supplemented with 1% glycerol, 0.05% tween80 and 10% Middlebrook OADC (oleic acid, dextrose, catalase, and bovine albumin). To select for mEmerald GFP-expressing Mab, zeocin (Invivogen) was included at a final concentration of 5μg/mL.

### Animal experiments

All cell isolation involving live mice was performed in accordance with the recommendations from the Guide for the Care and Use of Laboratory Animals of the National Institutes of Health and the Office of Laboratory Animal Welfare. Mouse studies were performed using protocols approved by the Institutional Animal Care and Use Committee (IACUC). All mice were housed and bred under specific pathogen-free conditions and in accordance with Michigan State University (PROTO202200127) IACUC guidelines. All mice were monitored and weighed regularly. C57BL6/J mice (# 000664) and Nrf2-/-mice (# 017009) were purchased from The Jackson Laboratory.

### BMDM and FLAM cell isolation, maintenance, and culture conditions

Primary BMDMs were generated by isolating marrow from femurs and plating in RPMI containing 25ng/ml M-CSF (Peprotech) and 10% FBS for 8-10 days. Cells were then washed with PBS and lifted with PBS-EDTA prior to plating for experiments. In indicated experiments, HoxB8-conditionally immortalized macrophages were used. These cells were isolated from C57BL6/J mice and were maintained in media containing 30ng/mL recombinant mGM-CSF (Peprotech), 10% fetal bovine serum (FBS) and 0.5μM β-Estradiol as previously described [59–61]. To generate BMDMs, cells were washed in PBS to remove estradiol, then plated in DMEM containing 25ng/ml M-CSF (Peprotech) and 10% FBS. Five to seven days later, cells were plated for experiments as described in figure legends. For FLAMs, C57BL6/J pregnant dam mice were euthanized by CO_2_ prior to cervical dislocation and fetal liver derived cells were obtained as previously described [38]. Cells were cultured in complete Roswell Park Memorial Institute medium (RPMI; Thermo Fisher) containing 10% FBS, 30ng/ml recombinant mGM-CSF (Peprotech), and 20ng/mL recombinant hTGF-β1 (Peprotech). All macrophages were incubated in 5% CO_2_ at 37°C. FLAMs were monitored regularly by flow cytometry for the continued expression of alveolar macrophage markers.

### Macrophage infections and assays

#### Mab infection of macrophage monolayers

Single cell Mab suspensions were prepared by resuspending logarithmic phase bacteria in the appropriate macrophage cell culture media followed by a soft spin at 58 xg to pellet large bacterial clumps. Macrophages were seeded at 5×10^5^/well in a 12-well and single cell Mab supernatants were used for macrophage spinfections (spun at 58xg for 5 minutes) at the indicated MOIs. Four hours later, infection media was removed and media containing 64μg/mL Amikacin was added for the remainder of the experiment. At each indicated timepoint (4 to 48 hours), macrophages were washed with PBS, lifted from plates by scrapping or Accutase^TM^ (BioLegend) treatment, then fixed in 4% paraformaldehyde. Infected macrophages were quantified using the BD LSR II (BD Biosciences) or Attune CytPix Flow Cytometer (Thermo Fisher Scientific) at the Michigan State University Flow Cytometry Core. Live and single macrophages were identified using forward and side scatter and the mean fluorescent intensity of infected cells was determined by the fluorescence in the GFP channel. All experiments included uninfected and unstained controls to set gates for infected macrophage quantification. Analysis was performed using FlowJo V10.

#### Mab infection of IFNγ activated macrophages

For IFNγ activated macrophage experiments, cells were treated with 25ng/mL IFNγ (Peprotech) for 16-20 hours. Following macrophage activation, IFNγ containing media was removed and infection media was added as described above. Four hours later, infection media was removed and media containing 64μg/mL Amikacin and 25ng/mL IFNγ was added for the remainder of the experiment.

#### Quantification of Mab intracellular growth

For intracellular growth experiments, macrophages were lysed at the indicated time points (12, 24 and 48 hours) with sterile cold distilled water. Following 10-fold serial dilutions on Middlebrook 7H10 agar supplemented with Middlebrook OADC and 5 μg/mL zeocin, samples were plated to perform colony forming unit (CFU) counts. Colonies were enumerated after 4-5 days of incubation at 37°C.

#### Quantification of Cell death

For flow cytometry cell death experiments, macrophages were lifted at the indicated time points (4 to 48 hours), then stained with Zombie Red Live/Dead stain (Biolegend). Once stained, cells were washed with PBS and fixed in 4% paraformaldehyde before being analyzed by Flow cytometry. For CellTiter Glo Cells were seeded in 96-well opaque plates the day prior to the experiment and stimulated with 25ng/mL of IFNγ (Prepotech) overnight. The following day, macrophages were infected as described above. At the indicated time points, viability measurements were performed following CellTiter Glo manufacture instructions. Briefly, 100μL of CellTiter Glo® Reagent was added directly to each well and plates were incubated for two minutes with shaking to induce cell lysis. Plates were then incubated for 10 more minutes at room temperature to stabilize luminescent signal and luminescence was measured on a Spark® multimode microplate reader (Tecan). Luminescence signal was normalized to uninfected cells for each condition.

#### Inhibitor/Activator experiments

For experiments blocking NF-kB the inhibitor BAY 11-7082 (Invivogen) was resuspended in ethanol and diluted in media to a final concentration of 5μM for experiments. For experiments activating HIF-1a using DMOG (Sigma-Aldrich) was resuspended in DMSO and diluted to a final concentration of 250μM in media for experiments.

### Griess Assay

The quantity of nitric oxide production by macrophages was determined by measuring its stable end product nitrite (Griess Assay, Promega). Briefly, 50uL of supernatant was transferred to a 96-well plate, followed by 50μL of sulphanilamide and 50μL of N-1-napthylethylenediamine dihydrochloride (NED) under acidic conditions. Following incubation, absorbance was measured at 540nm on a Tecan Spark 20M plate reader and nitrite concentrations were calculated using a standard nitrite curve per manufacture instructions.

### RNA isolation and quantitative RT-PCR

At the indicated time points (6 and 24 hours) macrophages from 6-well plates were resuspended in 1000 μL of TRIzol reagent (Life Technologies) and incubated for 5 minutes at room temperature. 200μL of chloroform was added to the homogenate, vortexed and centrifuged at 10,000 x g for 18 min at 4°C to separate nucleic acids. The upper aqueous phase was removed and combined with equal parts ethanol. This mixture was placed into a collection tube and protocols provided by the Zymo Research Direct-zol RNA extraction kit were followed. Quantity and purity of the RNA were checked using a NanoDrop and diluted to 5 ng/mL in nuclease-free water. The one-step Syber Green RT-PCR kit (Qiagen) reagents were used to amplify RNA according to manufacture instructions. Amplifications were monitored using the QuantStudio3 (ThermoFisher). Relative mRNA expression levels were calculated after normalization to β-actin.

### Cytokine analysis

Where indicated, supernatants were filter sterilized through a .2-micron filter before cytokines were quantified by a Luminex multiplex assay (Eve Technology). In addition, filter sterilized supernatants from Mab-challenged and control macrophages were harvested and the indicated cytokine proteins levels were determined using CXCL10, TNF or IL1α DuoSet ELISA kits (R&D Systems) following manufacture instructions. Absorbance (450nm) was detected on a Spark® multimode microplate reader (Tecan).

### RNA Sequencing and Analysis

This work was supported in part by Michigan State University through computational resources provided by the Institute for Cyber-Enabled Research. RNASeq read quality assessment, mapping, and counting were performed using a custom pipeline built in Snakemake version 7.32.4 (https://github.com/kaylaconner/olivelab-rnaseq/tree/main) [62]. Read quality was assessed using FastQC version 0.12.1 [63]. Read mapping was performed against the GRCm39 mouse reference genome using Bowtie2 version 2.5.1 [64]. Aligned reads counts were assessed using the featurecounts function from the Subread package version 2.0.6 [65]. Differential gene expression analysis was conducted using the DESeq2 package version 1.42.0 in R version 4.3.2 [66]. Pre-filtering was performed to keep only genes that had >10 counts in 3 or more samples.

Principal component analyses (PCA) were performed using the prcomp function with scaling on normalized count matrices of all genes in R version 4.3.2, and PCA visualization was done using the autoplot function in ggplot2 version 3.4.4 [67]. Magnitude/Amplitude (MA) plots were produced by plotting Log2(Fold Change) and Log2(Base Mean) values from DESeq2 in ggplot2 version 3.4.4 [67]. Heatmaps were produced in GraphPad Prism version 9.4.1 using normalized counts data from DESeq2. Venn diagrams were produced using the VennDiagram package version 1.7.3 in R version 4.3.2 [68]. Gene clustering analysis was performed using Clust 1.18.0 [69]. All gene ontology analysis was performed using the g:OSt functional profiling tool from the g:Profiler web server version *e111_eg58_p18_30541362* (database last updated on 25/01/2024) [70].

### Statistical analysis and data visualization

Statistical analysis was performed using Prism Version 10 (GraphPad) as indicated in the Figure Legends. Data are presented, unless otherwise indicated, as mean ± standard deviation. One-way or two-way ANOVA followed by Tukey’s post hoc test was used to identify significant differences between multiple groups, and Student’s t-tests were used to compare 2 groups.

## Acknowledgements

This work was supported by grants from the National Institutes of Health: R35 GM146795 (A.J.O.) The Attune CytPix, located in the MSU Flow Cytometry Core Facility, is supported by the Equipment Grants Program, award #2022-70410-38419, from the U.S. Department of Agriculture (USDA), National Institute of Food and Agriculture (NIFA).

